# A Novel, Open-Source Virtual Reality Platform for Dendritic Spine Analysis

**DOI:** 10.1101/2024.02.02.578597

**Authors:** Marike L Reimer, Sierra D Kauer, Curtis A Benson, Jared F King, Siraj Patwa, Sarah Feng, Maile A Estacion, Lakshmi Bangalore, Stephen G Waxman, Andrew M Tan

## Abstract

Neuroanatomy is fundamental to understanding the nervous system, particularly dendritic spines, which are vital for synaptic transmission and change in response to injury or disease. Advancements in imaging have allowed for detailed 3D visualization of these structures. However, existing tools for analyzing dendritic spine morphology are limited. To address this, we developed an open-source, virtual reality (VR) Structural Analysis Software Ecosystem (coined “VR-SASE”) that offers a powerful, intuitive approach for analyzing dendritic spines. Our validation process confirmed the method’s superior accuracy, outperforming recognized gold standard neural reconstruction techniques. Importantly, the VR-SASE workflow automatically calculates key morphological metrics such as dendritic spine length, volume, and surface area, and reliably replicates established datasets from published dendritic spine studies. By integrating the Neurodata Without Borders (NWB) data standard and DataJoint, VR-SASE also aligns with FAIR principles--guidelines aimed at improving the findability, accessibility, interoperability, and reusability of digital assets--enhancing data usability and longevity in neuroscience research.

**Motivation:** Technological limitations of available image-analysis tools for analyzing 3D fine-structure hinders effective research and is often costly. An accessible and efficient solution is crucial to overcome these research challenges. We addressed this by integrating the NWB data standard and DataJoint technology into an open-source, virtual reality workflow, enhancing dendritic spine analysis.

## Introduction

Dendritic spines are microscopic structures that serve as morphological sites of synaptic contact between neurons in the brain and spinal cord. Importantly, dendritic spine morphology directly influences how electrical inputs through synapses are received, transduced, and ultimately processed by the postsynaptic neuron^1^. As such, dendritic spine structure plays a critical role in supporting synaptic and circuit functions, making them a vital visual proxy for understanding the workings of the nervous system^2^. In pathology, abnormal dendritic spine structures (termed “dendritic spine dysgenesis”) is observed in multiple neurological and psychiatric disorders including autism, schizophrenia, Huntington’s Disease, chronic pain, and Alzheimer’s Disease^3, 4^.

Given the significance of dendritic spine morphology in understanding neural circuit function and pathology, it is crucial to have efficient methods for accurately measuring this anatomy. Historically, studies have primarily used two-dimensional (2D) image analysis. This is a process that necessitates manual or semi-automatic pixel-based tracing and extrapolation of dendritic spines in two dimensions, which are then projected or extrapolated to interpret observations within the real third dimension. However, these traditional approaches are not only tedious and time-consuming but also prone to systematic assumption and human subjective error. This is largely due to these tools’ dependence on extrapolating 3D metadata from 2D image datasets collected by investigators. Overall, while tools offer valuable insights and have paved the way for advancements, there is still a need for improved accessibility, usability, and comprehensive 3D analysis capabilities.

Here, we developed a Virtual Reality Structural Analysis Software Ecosystem (VR-SASE). The main features of VR-SASE are its accessible user interface, standardized data, automated data extraction and integration capabilities, and low cost. Users engage VR-SASE through workflow (shown in Figure 1), first, rendering the neuronal model in VR space based on microscopy-captured images, and then segments structures for analysis, e.g., dendritic spines, from the 3D model using virtual “slicing” discs. Software automates the remainder of segmentation and data management. Finally, in contrast to commercially available software packages, e.g., Neurolucida and Imaris, that require ongoing licensing contracts, VR-SASE is open-source, without recurring expense, making it an ideal option for research teams with limited budgets.

**Figure 1A:**
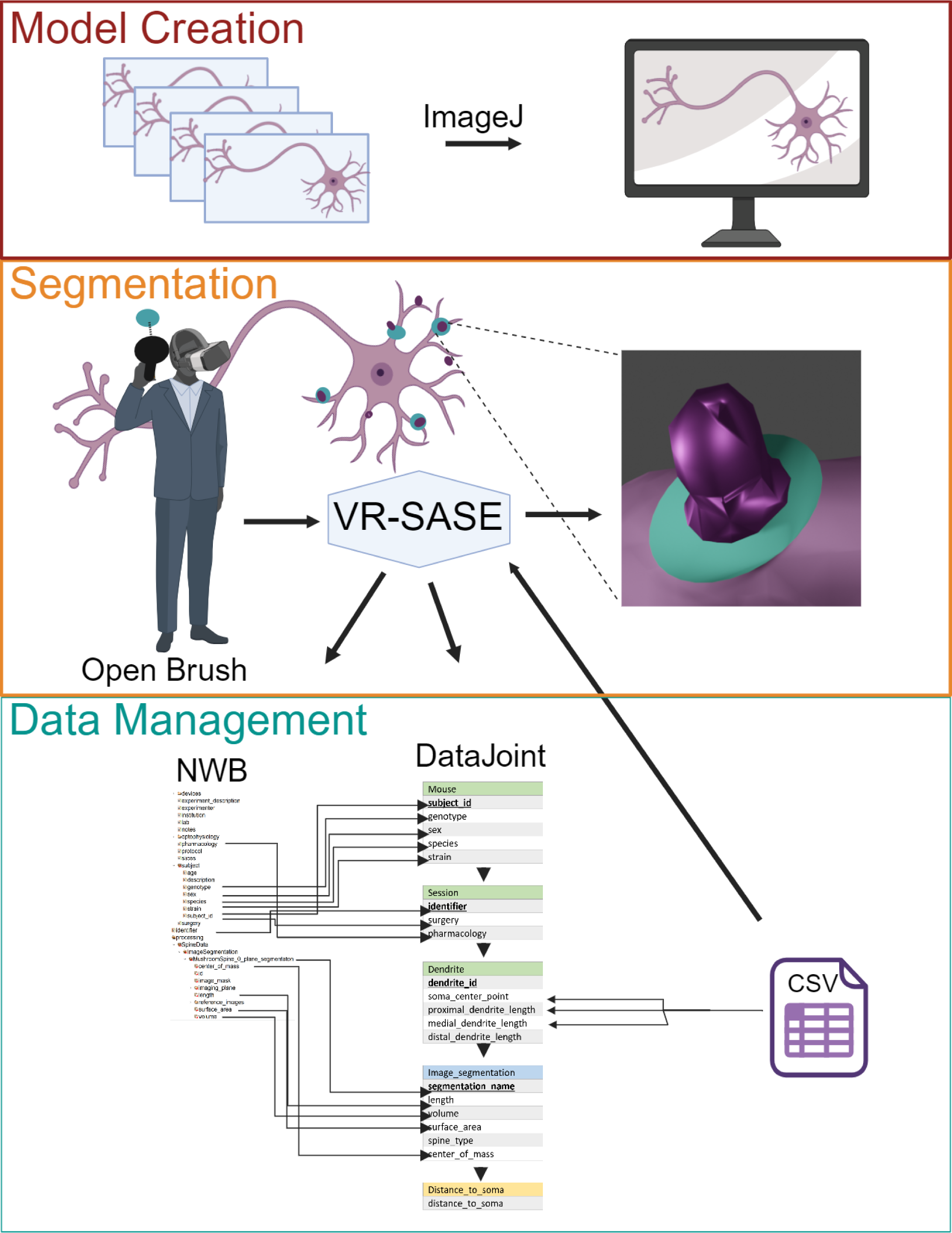
The VR-SASE Workflow consists of three phases: 1) Model Creation: The top panel shows neural tissue converted to a 3D model in the .obj format using ImageJ. 2) Segment Dendritic Spines: The model neuron is loaded into the Open Brush VR environment where the users place discs to “slice” off spine segments. The model is exported and opened in Blender where a addon completes the segmentation. 3) Data Management: VR-SASE automates saving morphological data in NWB files, integrates additional dendrite measures from a .csv file, unifying data in DataJoint for quality control and advanced analytics.

We assessed the utility of VR-SASE using two approaches: First, we compared the platform tool against conventional imaging methods, noting its ability to match their results while providing additional metrics like dendritic spine surface area and volume. Second, we benchmarked the VR-SASE toolset use against the DIADEM challenge (DIgital reconstruction of Axonal and DEndritic Morphology)^5^. VR-SASE outperformed the DIADEM model in terms of accuracy and may provide better utility than existing methods for analyzing dendritic spines. The open-source framework of VR-SASE fosters collaborative and community-led development, offering researchers a straightforward and accessible means to comply with FAIR data principles, enhancing data findability, accessibility, interoperability, and reusability. Finally, VR-SASE transforms labor-intensive 3D image analysis procedures into a more intuitive, precise, and efficient workflow, setting the stage for deeper insights into neurological and psychiatric disorders.

## Results

### Neurolucida, ImageJ, and VR-SASE comparison highlights advances in visualization, accuracy, and precision

Figure 2 provides a visual comparison of the Neurolucida, ImageJ, and VR-SASE interfaces. A representative neuron traced in Neurolucida is depicted in (Fig. 2A), an enlargement shows individual spines marked with thin red lines in (Fig. 2B). (Fig. 2C) shows the same spines visualized in VR-SASE. (Fig. 2D) shows the leftmost spine from (Fig. 2B) in ImageJ, maximally enlarged and processed to enhance visibility. Red lines, placed by four experts, capture the length of the spine, demonstrating the inherent variability of manual measurements. In contrast, the same spine is shown from the VR-SASE workflow, (Fig. 2E), with greatly enhanced resolution. The spine tip is clearly marked by an orange sphere (indicated by an arrow), minimizing ambiguity in length measurements. Overall, VR-SASE offers superior visualization with increased accuracy and precision.

**Figure 2:**
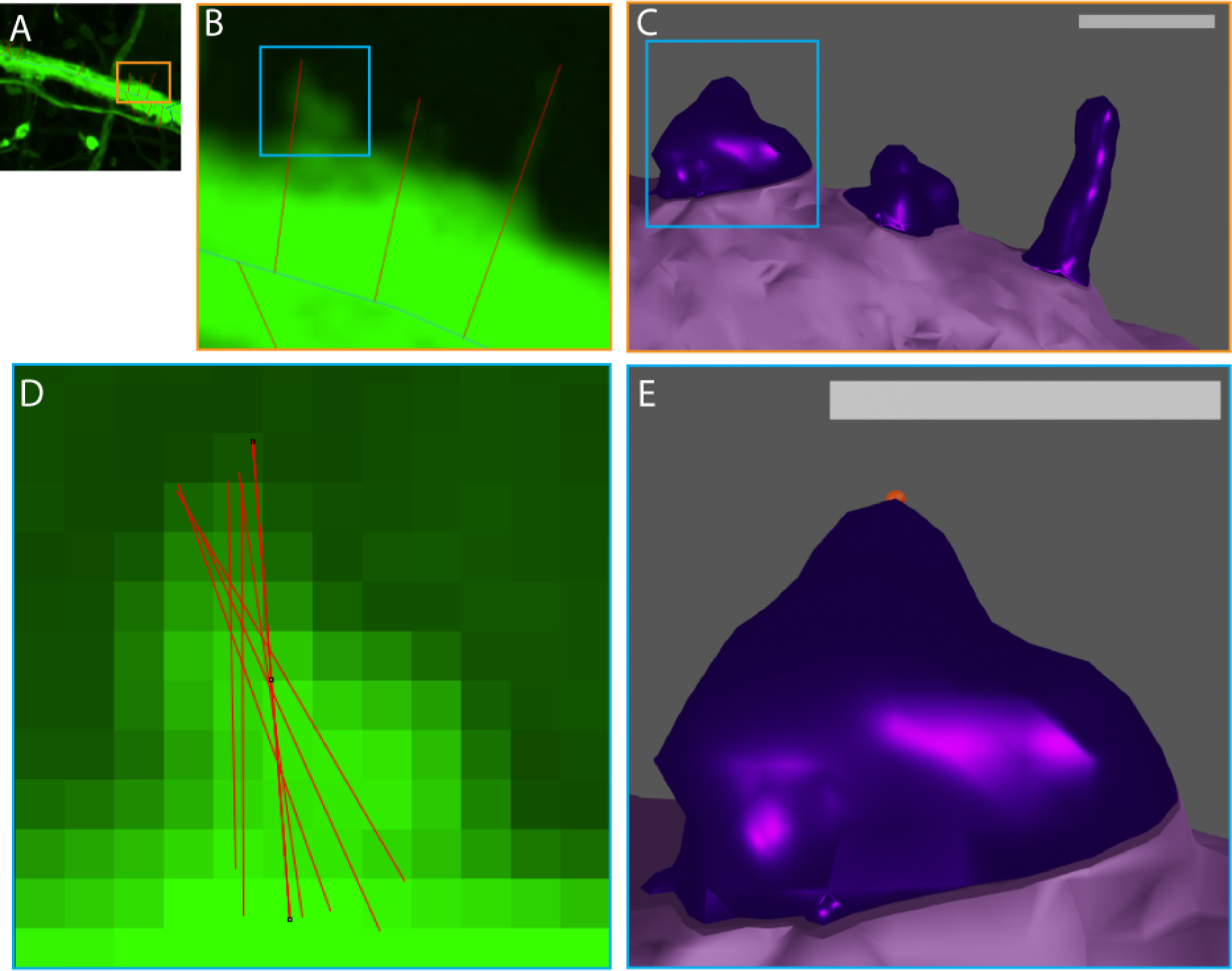
Neurolucida, ImageJ, and VR-SASE comparison highlights advances in visualization, accuracy, and precision. (Fig. 2A) shows a neuron traced in Neurolucida. (Fig. 2B) is an enlargment showing spines as an overlay marked with thin red lines denoting spines. (Fig. 2C) shows these spines in VR-SASE, with much greater clarity and detail. (Fig. 2D) shows the leftmost spine, maximally enlarged. Four expert analysts placed red lines denoting length, indicating the high degree of variability inherent in manual measurements. (Fig. 2E) is an enlargment of the left-most spine shown in (Fig. 2B,D). The spine tip (arrow) is marked with a sphere, to aid in visualization, highlighting how VR-SASE decreases abiguity in length measurements.

### Abnormal dendritic spine density increased in the proximal region of the dendrite

To validate VR-SASE, we analyzed spinal cord neuronal data presented by Dr. Kauer et al. at the 2023 Society for Neuroscience poster presentation: Pak1 inhibition with Romidepsin attenuates H-reflex excitability after spinal cord injury, and at the 2022 Paralyzed Veteran’s Association in Dallas Texas.

In this study, blinded investigators performed a Sholl’s analysis with Neurolucida to examine changes in the distribution of dendritic spines in three regions around the neuron’s soma: proximal (0-30 µm), medial (30-60 µm), and distal (60-90 µm). Dendritic spines were traced on sequential images spanning each dendrite using Neurolucida’s ‘stack scrolling’ interface. This method requires switching between images to mark the endpoints of the spine.

Using Neurolucida, the SCI group had a higher proximal total spine density: Unpaired T-Test: Sham (M = 0.1513) and Injured (M = 0.3734); p = 0.0058, t=3.041, df=23, SEM =0.07303. (Fig. 3A), thin spine density: Unpaired T-Test: Sham (M = 0.1321) and Injured (M = 0.2617); p= 0.0322, t=2.280, df=23, SEM =0.05681. (Fig. 3B) and mushroom spine density: Mann Whitney test: Sham (M = 0, n=9) and Injured (0.06399, n=16); p= 0.0356, U = 35.50. (Fig. 3C).

**Figure 3:**
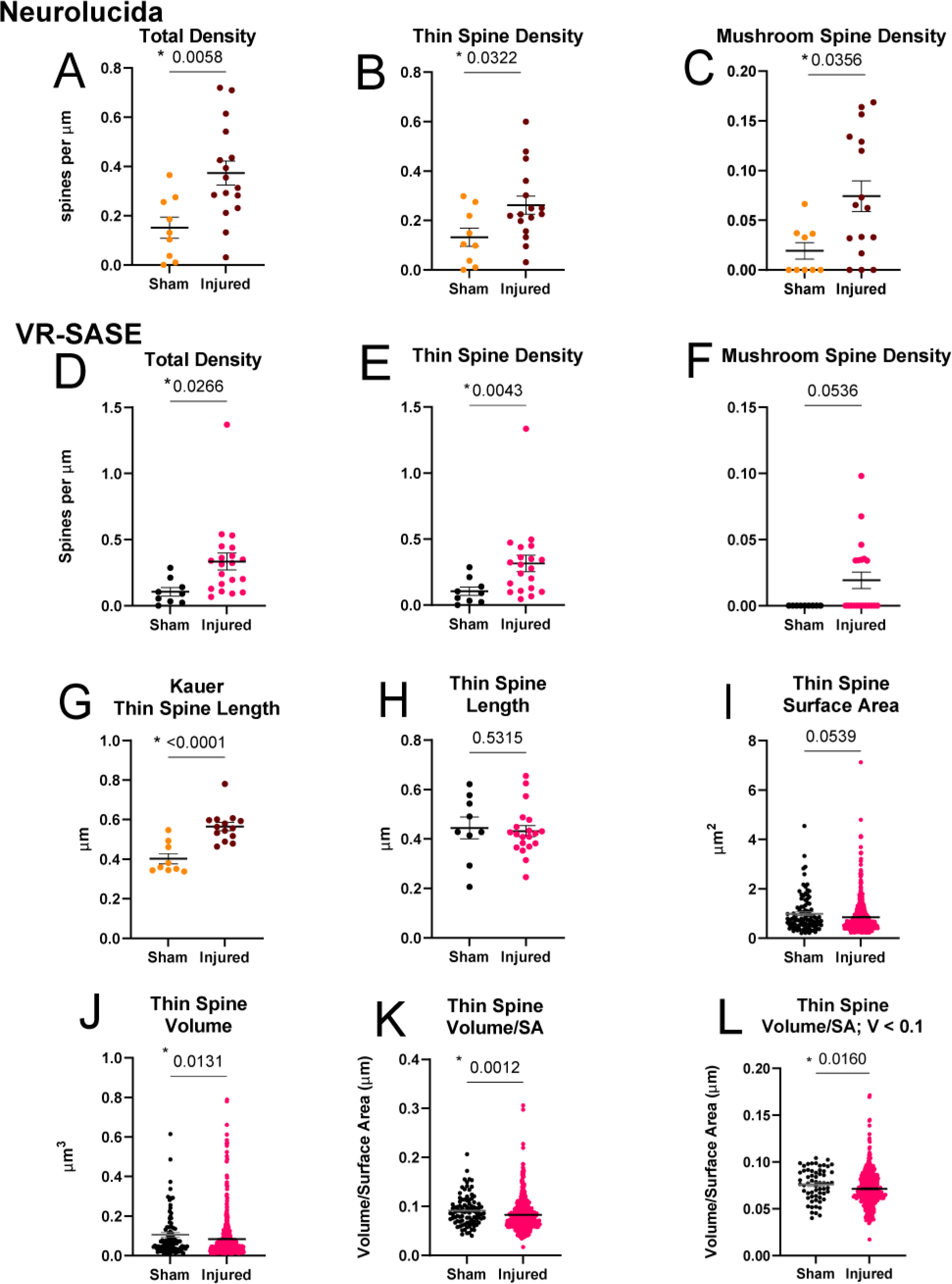
SCI Increases Proximal Dendritic Spine Density and Abnormal Morphology. VR-SASE recreated the Neurolucida-traced findings of Kauer et al 2022, 2023. Their analysis showed that compared to Sham, the SCI group had a significantly higher proximal total spine density (Fig. 3A), thin spine density, (Fig. 3B), and mushroom spine density (Fig. 3C). We likewise found in the SCI group significantly higher proximal total spine density (Fig. 3D) and thin spine density, Fig 3E; mushroom spine density approached significance, as shown in Fig 3F. Kauer et al found that SCI mice had significantly longer thin dendritic spines than Sham (Fig. 3G) using ImageJ, however VR-SASE did not show a significant difference between groups (Fig. 3H). Using the additional metrics provided by VR-SASE, we assesed thin spine Surface Area (Fig. 2I) and Volume (Fig. 3J). No differences were found between groups for Surface Area; however injured mice had dendritic spines with less volume. We assessed the Volume to Surface Area of thin spines and found that the injured group had decreased values (Fig. 3K). This was not true of spines greater than 0.1 µm^3^ but was observed in dendritic spines smaller than 0.1 µm^3^ (Fig. 3L).

Moving freely in a VR space, VR-SASE enabled users to visualize and segment neuroanatomy from any perspective. Using our pipeline, we analyzed images from the Kauer et al., 2023, 2022 dataset. Our spine density findings (Fig. 3 D-F) are largely in line with Kauer et al. 2023, 2022 results. We focused our analysis on the proximal region and likewise found an increase in total spine density: Mann Whitney test: Sham (M = 0.09019, n=9) and Injured (M = 0.3318, n = 20); p= 0.0017, U = 26. (Fig. 3D). We also observed the same trend for thin spine density: Mann Whitney test: Sham (M = 0.09019, n=9) and Injured (0.09019, n=9); p=0.0043, U = 31 (Fig. 3E). We found mushroom spine density that approached significance: Mann Whitney test: Sham (M = 0, n = 9) and Injured (M = 0, n = 20); p = 0.0536, U = 54 (Fig. 3F). These findings are in line with previous results showing that SCI induces an abnormal increase in dendritic spine density ^2, 6–8^.

### SCI induces abnormal dendritic spine morphology

Changes in the size and shape of dendritic spines can alter how signals are transmitted and how efficiently synapses function, impacting the physiology of neurons.^2^ Following precedent set by our group^6^, Kauer et al. 2023, 2022 measured spine length manually using FIJI, tabulating values in Excel. These findings showed that SCI led to an increase in the total length of dendritic spines, compared to the sham, driven largely by an increase in thin spine length: Unpaired T-Test: Sham (M = 0.4020) and Injured (M = 0.5647); p < 0.0001, t=4.893, df=21, SEM = 0.3325 (Fig 3G). This is in agreement with our published work^6^ and is a general trend in neurons after SCI ^8, 9^. We did not find that the length of dendritic spines increased following SCI: Mann Whitney test: Sham (M = 0.4283, n=9) and Injured (M = 0.4206, n=20); p= 0.5315, U = 76 (Fig 3H).

We next sought to leverage the additional measures provided by VR SASE to quantify how contusion injury induces abnormal morphological changes in dendritic spines. We did not find any significant differences between groups for Thin Spine Surface Area (Fig 3I). However, we found that thin spine volumes were smaller in the injured animals: Mann Whitney test: Sham (M = 0.06853, n=94) and Injured (M = 0.04969, n=526); p= 0.0131, U = 20759 (Fig 3J). We also found a significant difference between the Volume to Surface Area ratio between injured and uninjured animals. The injured group had a significantly lower ratio: Mann Whitney test: Sham (M = 0.08784, n=94) and Injured (M = 0.07661, n=526); p = 0.0012, U = 19562 (Fig 3K).

We removed all spines less than 0.1µm^3^ and found that this effect was eliminated (not shown). By restricting our analysis to spines with volume less than 0.1µm^3^, we confirmed that this effect was due to the tiny spines: Mann Whitney test: Sham (M = 0.07719, n=61) and Injured (M = 0.06928, n=409); p = 0.0160, U = 10096 (Fig 3L).

The time required to generate these analyses was substantially decreased when compared to the same analyses carried out in Neurolucida and ImageJ. Positioning the disc “slicers” on VR neurons took an average of 17 minutes per neuron (Extended Data), representing a substantial decrease in researcher effort. By augmenting length and density measures with surface area and volume, VR-SASE enhances our understanding of how structural changes in dendritic spines contribute to pathological states. By providing these measures efficiently, VR-SASE accelerates and expands our research capabilities.

### VR-SASE validated with the DIADEM gold standard

The DIADEM challenge was a widely lauded effort consisting of elite research groups pitting automated neural reconstruction algorithms against each other, generating neural models long considered to be the gold standard^10^. To assess the accuracy of our VR segmentation, we reconstructed a drosophila olfactory neuron, OP-09, from the DIADEM final round. We then compared our reconstructed neuron (Fig. 4B) with the gold standard reconstruction (Fig. 4C) by super-imposing them on the maximum projection of OP-09 (Fig. 4A). (Fig. 4D) shows our reconstruction of OP-09 on top of the maximum projection. Deviations from true morphology are visible as green around the edges. The gold standard reconstruction (Fig. 4E) has 58% more green visible, indicating that our tracing is more accurate. (Extended Data) Our segmentation of OP-09 is shown in Fig. 4F. Fig. 4G is an enlargement showing the placement of the teal ‘slicer’ discs. VR-SASE used the teal discs to “slice” the complex neuron into its component pieces. Each section was assigned a random color, highlighting segmentation accuracy, as well as Blender’s utility in visualizing information.

**Figure 4:**
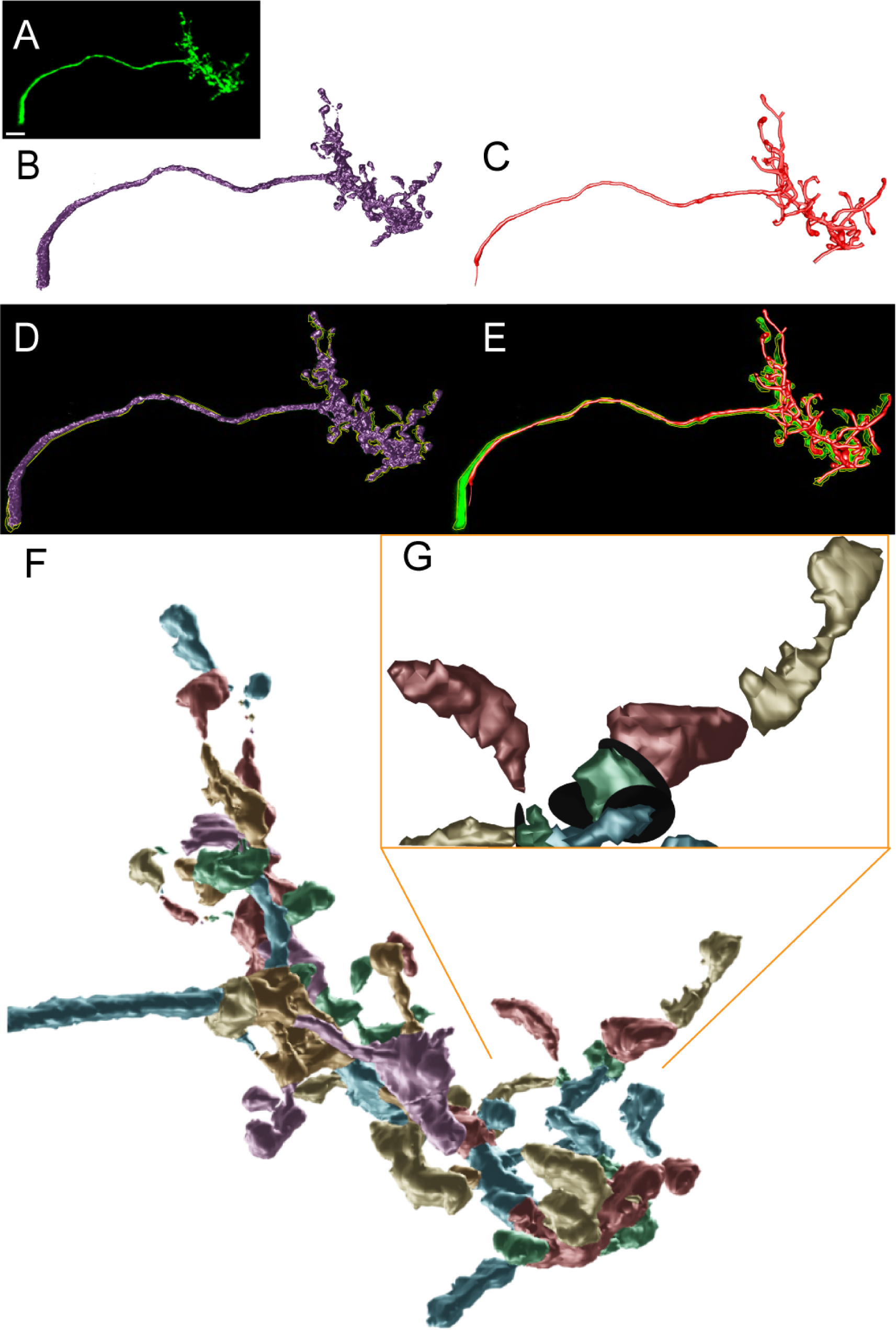
VR-SASE Validated with the DIADEM Gold Standard (Fig. 4A) shows OP-09, a neuron from the DIADEM challenge. Scale bar is 10 µm. (Fig. 4B) shows the VR-SASE reconstruction and (Fig. 4C) shows the gold standard reconstruction. We super-imposed the reconstructions (Fig. 4D and E), respectively, on the maximum projection (Fig. 4A). We quantified the green regions and found the gold standard reconstruction had 58% more green visible, indicating that VR-SASE depicted anatomy more accurately. (Fig. 4F) shows results of segmentation indicated with different colors. The expanded view in (Fig. 4G) is an enlargement demonstrating how black ‘Slicer’ discs were placed to segment the neuron.

We examined the model in Blender, using its native 3D-Printing addon and found the volume to be 1216.1861 µm^3^ and the surface area to be 3061.1194 µm^2^. The time to position the teal discs “slicers” was just under 15 minutes. The benchmark for manual reconstructions of simple neurons is approximately 20 minutes, although complex neurons such as OP-09, take longer. ^5^

By reconstructing OP-09, a gold standard neuron with VR-SASE, we demonstrated that the accuracy of VR-SASE surpassed the gold standard by 58% (Extended data). Additionally, VR-SASE provided additional meta-information about the neuron in its ability to generate insights with volume and surface area data.

### Advanced analytics with VR-SASE

In Pchitskaya et al., investigators employed a new method to aid in dendritic spine classification by creating a chord length distribution histogram (CLDH)^11^. This technique measures the internal chords of dendritic spines, creating a histogram which can be used to distinguish between dendritic spine clusters. Fig. 5A shows a dendritic spine. In Fig. 5B, the surface of the spine has been removed to show its internal cords. Pchitskaya generated their cords randomly, which could lead to under-sampled regions. To ensure complete coverage, we created cords at every vertex in the model. We excluded nearest neighbor vertices and sampled one in every ten of their possible connections, generating a rich collection of cords. Fig. 5C depicts the CLDH for the dendritic spine. This underscores how data standardization enabled VR-SASE to employ analysis techniques.

**Figure 5:**
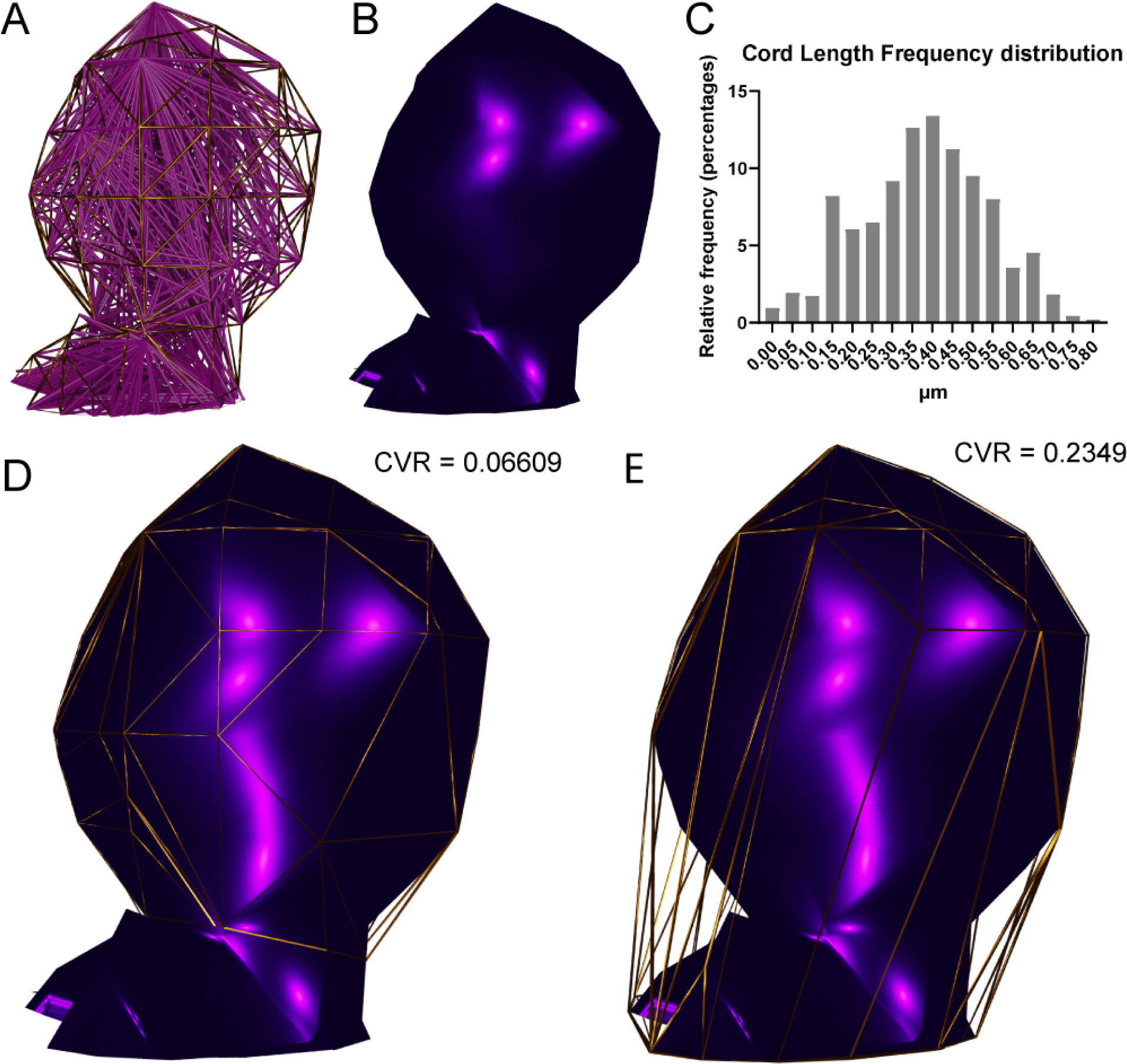
VR-SASE post-hoc analyses demonstrate advanced analytic capabilities. (Fig. 5A) shows a dendritic spine. (Fig. 5B) shows the interior of the spine showing cords created with a post-hoc analysis in VR-SASE. (Fig. 5C) shows the CLDH, a distribution of their lengths. Ptichkaya et al. demonstrated that this technique improves classification accuracy^16^. Their proposed method randomly creates cords in the interior of the spine, which introduces inaccuracies by over and under sampling regions. We mitigate these errors by creating cords at each vertex in the 3D model of the spine. (Fig. 5D) depicts our post-hoc analysis for Concave Volume Index. The difference in volume between a spine head and its convex hull, which is a shell around the spine without concave surfaces. This method was pioneered by Kashiwaki et al. to assess dendritic spine head morphology.^12^ Given that strict classification of dendritic spines relies on the presence of an indent (concave surface)^27^ this measure has utility in dendritic spine clusterization and could be augmented still further with DataJoint advanced analytics.

Next, we sought to expand VR-SASE’s capabilities by calculating the CVI for a dendritic spine. Fig. 5D shows the dendritic spine from Fig. 5A with its head surrounded by a convex hull (gold mesh) recreating the method used by ^12^. Fig. 5E shows this method used quantify the concavity of the entire dendritic spine. Applying this measure of dendritic spine concavity to DataJoint queries, as we did with dendritic spine length and volume, can foster innovative dendritic spine clustering techniques.

## Discussion

Dendritic spines are neuronal elements that form the structural basis for synaptic contact between neurons of the central nervous system. Their morphology dynamically changes in response to both environmental influences and internal factors, under both normal and pathological conditions. Historically, analysis of these structures has been hindered by the limitations of 2D imaging methods. VR-SASE offers an open-source, e.g., more affordable, intuitive, accurate, and efficient workflow compared to traditional methods and commercial software.

The DIADEM challenge aimed to accelerate automated neural tracing but fell short of its target^13^. Big Neuron, the next generation platform for validating neural reconstructions algorithms was recently released ^10^. However, neither addresses dendritic spines. The need for improved analysis tools led to commercial solutions like Imaris and Neurolucida, which are costly to maintain. Accessing Neurolucida data requires a proprietary workstation with an active license. Unlike VR-SASE, automated solutions offered by these platforms do not allow for manual refinements and do not offer the versatile morphological measures provided by ImageJ, which was developed by the NIH^14^.

The need for open-source dendritic spine analysis created an abundance of analytical software platforms, which have been reviewed elsewhere^15, 16^. Briefly, 3dSpAn^17^, DXplorer ^18^, and Spine Tool ^11^ are recent and remarkable advancements for analyzing dendritic spine morphometries, however, they present obstacles for effective analysis. Their reliance on 2D interfaces (e.g. flat computer screens) limits visualization, manipulation, and segmentation of 3D objects.

In this study, we introduced VR-SASE that leverages VR technology to address these challenges and facilitates dendritic spine analysis. VR-SASE replaces the complex computations underlying existing platforms by simply placing virtual slicing discs to create segments. Moreover, VR-SASE provides users with excellent model visualization in a room-scale VR environment and precise control through its intuitive VR environment and semi-automated segmentation. The immersive VR environment overcomes the limitations of traditional 2D interfaces enabling researchers to visualize minute dendritic spine morphology from all perspectives. Finally, the VR-SASE workflow automatically generates key morphological measures, including dendritic spine length, volume, and surface area. These measures play crucial roles in understanding electrical interactions,^19, 20^ synaptic strength^21^, and calcium dynamics^22, 23^. Automated collection of morphological data empowers VR-SASE users to interpret their anatomical data in less time with less subjective error risk. Integration with NWB and DataJoint further enables the VR-SASE workflow to follow FAIR data compliance^24^, improving data interoperability, streamlining data organization, and ensuring dataset completeness and minimizing human errors.^25^ Furthermore, files created by VR-SASE and shared via the free online NWB hosting site, DANDI Archive, can satisfy the requirements of FAIR compliance^24^.

We applied VR-SASE to the investigation of dendritic spine changes following SCI, demonstrating its accuracy and utility by recreating Kauer et al. findings that SCI causes abnormal dendritic spine density. We extended Kauer et al. 2022, 2023 findings by showing that SCI causes abnormally decreased thin spine volumes with significantly smaller volume-to-surface-area ratios^22, 23,26,27^. In this vein, while our 0.2 µm minimum length criteria includes fine structural resolution, included structures may not fall within identity criteria for dendritic spines in other tissue types or analytical approaches^28^. Nonetheless, our finding of decreased volume-to-surface-area ratio in tiny spines of injured mice demonstrates VR-SASE’s capabilities in measuring this important metric.

We further validated the accuracy of our method by reconstructing a drosophila olfactory neuron from the DIADEM challenge and our results compared favorably against the gold standard reconstruction.

Our reconstruction depicted the neuroanatomy more accurately, demonstrating the effectiveness of VR-SASE in creating extremely accurate models. We then demonstrated the utility of data standardization and the adaptability of VR-SASE by conducting two *post hoc* analyses. First, by creating a CLDH with improved cord distribution, we expanded on the methodology proposed by Pchitskaya et al ^11^. Second, we calculated the CVI for a dendritic spine head, using the method published by Kashiwagi^12^. We calculated the CVI for the entire dendritic spine. Augmented with DataJoint’s analytics, this measure of dendritic spine concavity could refine collections of dendritic spines, just as our analysis of Kauer et al. 2022, 2023 selected only spines satisfying established length and volume parameters. Furthermore, our CVI data also demonstrated that VR-SASE can segment dendritic spine necks, a step towards quantifying this vital compartmental morphology^20^.

Blender is an optimal platform to integrate diverse modalities as it is already widely used in academic research. A recent Google Scholar query for “Blender Model” returned more than 200,000 results from diverse fields. The use of VR is making substantial inroads in the neuroanatomical studies ^13, 29–31^. Applying VR to analyze dendritic spines is relatively novel, and will continue to evolve. VR-SASE is amenable to machine learning approaches, such as PointNet, which would further expedite analyses^32^. Blender’s versatile Boolean modifiers can segment virtually any 3D shape, making VR-SASE an excellent hub for training AI on 3D models throughout biology.

Beyond the focus on dendritic spines, VR-SASE has the potential for a diverse biomedical research application. For example, the platform could analyze axonal morphology, neuronal circuitry, or even vital organ reconstructions through the body. Compatibility with electron microscopy workflows makes VR-SASE a broadly applicable dendritic spine analysis platform and facilitates interchange of 3D formats. Integration with the NEURON simulation environment also provides a route to *in silico* functional modeling.

An inherent strength of VR-SASE lies in its standardized data, enabling integration with existing tools. We implemented advanced DataJoint functions in a Jupyter notebook, so our advanced DataJoint analytics can be recreated from this file. DANDI Hub and Jupyter Lab facilitate Jupyter Notebook integrations with other programming languages including Julia, R, and Matlab. This interoperability across diverse programming languages contributes to a richer understanding of dendritic spine dysgenesis.

In conclusion, VR-SASE marks a substantial advancement in neuroanatomical research, especially in dendritic spine analysis. It surpasses traditional methods in accuracy, efficiency, and intuitive usage. VR-SASE’s application in anatomical studies and its adaptability for a wide range of biomedical research applications underscore its potential as a versatile tool in biology.

### Limitations of Study

The mushroom spine density we calculated is lower than Kauer et al. 2022, 2023, only approaching significance. This is likely due to the sensitivity of mushroom spines to thresholding, which my become detached due to their faint necks. We did not find that SCI causes an increase in spine length as Kauer et al. 2022, 2023 and others have ^2, 4^. Several factors may have contributed to this finding, including the length measurement variability within and between the VR-SASE and ImageJ, and variations arising from stubby spine morphology, a recognized challenge in the field ^18^. (See Figure S3). Another possibility is that modeling protocol requires greater refinement, as threshold settings impact the presence and connection of dendritic spines. Dendritic spine neck morphology strongly influences electrical behavior ^20^ so optimizing model creation is therefore of utmost importance to the use of VR-SASE. Using ImageJ’s gamma correction to enhance the contrast between dendritic spines and the background could address these issues ^33^.

Despite the advancements presented by VR-SASE, there are areas for further improvement. Integration with Python libraries such as Pandas would streamline analyses. Additionally, linking VR-SASE with the Cedar Metadata Workbench would optimize the metadata workflow [34], and automating the Sholl’s analysis would increase efficiency. Challenges arise from datasets at this scale, which push Blender to its limits. Manual intervention is occasionally necessary, prompting the creation of a dedicated “Manual Segmentation” tool (Fig 2C). While researcher involvement remains crucial, Blender’s hotkeys accelerate quality control. Future developments may capitalize on increasing computational power and Blender’s own VR environment to further reduce reliance on 2D interfaces.

The open-source nature of VR-SASE promotes collaboration and community-driven development. Researchers are encouraged to contribute to the platform’s improvement by sharing code, proposing features, and reporting issues through the Github repository. This collective effort fosters innovation, accelerating the advancement of dendritic spine analysis techniques. By mitigating the analysis bottleneck, VR-SASE plays a pivotal role in reducing the time required for developing new therapies and treatments. In conclusion, VR-SASE fills a critical gap in dendritic spine analysis by integrating VR technology into a standardized, open-source software application, marking a significant step forward in cell biology methods.

## Acknowledgements

The work is funded by grants from the Paralyzed Veterans of America (PVA) Research Foundation, the Department of Veterans Affairs (VA) Medical Research Service and Rehabilitation Research Service (RX003728-01A1) and The Taylor Foundation for Chronic Disease. and the Flaherty and Wilshire Walsh Endowments, as well as the SPiRE Program. The Center for Neuroscience and Regeneration Research is a Collaboration of the Paralyzed Veterans of America with Yale University. We thank Jennifer Carrara and Pamela Zwinger for their excellent technical assistance.

## Author Contributions

Marike L. Reimer: Software and methodology development, manuscript, figure creation, segmentation, and data analysis.

Sierra D. Kauer: Data generation, manuscript review, SCI subject matter expert.

Curtis F. Benson: Dendritic spine subject matter expert, project consultation.

Jared A. King: Technical review, figure creation.

Siraj Patwa: Software prototype development and VR segmentation.

Sarah Feng: VR segmentation prototype development, methods documentation.

Maile A. Estacion: Developed VR segmentation prototype.

Lakshmi Bangalore: Writing - review & editing.

Stephen G. Waxman: Review.

Andrew M. Tan: Developed vision, procured funding, supervision, writing.

## Declaration of Interest

The authors have no conflicting interests to declare.

### STAR Methods

### Image Analysis Workflow

Neuronal images were collected from mice that received either a contusion injury or a sham injury Kauer et al. 2022, 2023. To identify α-motor neurons, we employed a screening process based on data from previous studies ^6, 27^. These neurons, located in the ventral horn Rexed lamina IX, met the criteria of having a soma diameter larger than 25 μm and a cell body cross-sectional area greater than 450 μm². For further analysis, we included only YFP+ α-motor neurons with visible cell bodies and dendritic branches at least 45μm in length. Three-dimensional reconstructions of motor neurons were analyzed to examine dendritic spine density and distribution. Thin, stubby, and mushroom spines on each dendritic branch were identified and marked by placing a disc to “slice” or segment them apart from the dendrite.

Putative mushroom spines were assessed from orthogonal perspectives and classified as mushroom spines if their head was wider than their neck is long, which is an established classification ^4, 6, 9, 27^. Ambiguous cases were handled by comparing the virtual cursor with the length of the dendritic spine neck and the width of the dendritic spine head.

To calculate dendritic spine densities, we pooled all non-mushroom spines into the category “Thin” ^4, 6, 9, 27, 34^. Limiting our classification to “Thin” and “Mushroom” enabled us to measure subtle variations in spine distribution. Dim neurons and those with pronounced YFP puncta were excluded from the study as they produce digital models with high noise to signal ratios, obscuring potential dendritic spines.^35^

### ImageJ

We used version 1.535f1 of NIH ImageJ (https://imagej.nih.gov/ij/download.html) to process images and convert them to .obj format, which can be opened in Open Brush (https://openbrush.app/). To ensure high fidelity conversion, clear separation between dendritic spines and the background was necessary. To achieve this, ImageJ’s Autocorrect Brightness and Contrast function was applied at the center of the image stack. A border of each image was selected and filled using the rectangular selection tool, creating a bar of known dimensions to reconstruct scale. The image type was set to 8-bit and then saved as a pgm file before saving as in obj format. The resampling factor was set to 1 to prevent smoothing. Multiple threshold settings were tested for each image stack, and a region was compared with the original image stack to ensure the presence of all dendritic spines in the region.

### Segmentation with Open Brush

Open Brush, a room-scale virtual reality 3D painting platform was originally developed by Google under the name Tilt Brush but was renamed Open Brush after they released it as open source. Open Brush is available through multiple platforms, depending on VR hardware requirements. We used two VR headsets to demonstrate platform independence (see Hardware Descriptions). We installed Open Brush on the Oculus Quest 2 VR headset, produced by Meta Platforms, using the Meta Store interface. This interface is not available on the Oculus Rift S, so Open Brush software was downloaded through Steam, a digital distribution service.

The obj files created by ImageJ were placed into Open Brush’s media library, allowing import into the virtual environment, where a blinded researcher scaled them to the size of the room. The neural model was then “pinned” to ensure that the “slicer” discs remain attached to the model. Slicers are flattened cylinders created in Blender and imported from Open Brush’s media library. They were resized and positioned creating a clean interface when they are subtracted from the 3D mesh of the neuronal reconstruction at a later stage.

For tracing procedures, the main light source within the VR environment was positioned between the user and the neuron. This control over lighting provided clear visibility, good contrast resolution on the digitized neuronal tissue model, and consistency between models. Dendritic spines were segmentedd on one side of the dendrite, and then the user moved to the other side of the neuron, adjusting the light to shine directly on the neuron (as above to ensure clear visualization of the neuronal tissue model).

### Segmentation with Blender Addon

We developed a custom addon for Blender, an open-source 3D modelling suite, using their Application Programming Interface (API) for the Python programming language. See Fig S1. We used the CGFigures addon to install the PyNWB and DataJoint libraries to enable NWB file creation and database integration, respectively.

Open Brush allowed positioning Slicer discs on a neuronal model which can be exported as .fbx files. These were imported into Blender. The models and discs were then scaled to their original dimensions and aligned with Blender’s coordinate system. Slicers give their names to the dendritic spines they segment. This standardizes naming conventions, without restricting the scope of possible analyses.

To ensure data integrity, the original .obj file of the neuronal model was re-imported and the model from the .fbx file discarded, retaining only the correctly positioned Slicer discs from VR segmentation.

Segmentation with the VR-SASE Blender Addon begins with copying all the slicers and joining them into a single mesh with Blender’s ‘Join’ functionality. The merged slicers are then applied to the neural model as the object of a “Boolean Difference” modifier operation. This removes the portions of the neural model connecting the dendritic spines to dendrite. The “Separate Meshes” button on the VR-SASE Segmentation Tools Panel creates individual meshes of each spine.

The separated dendritic spines must have their origin updated through Blender’s Graphical User Interface (GUI). Then the ‘Slicer’ discs must be placed in a collection. To automatically segment the dendritic spines, they were selected along with the collection containing the Slicers.

The “Segment Solid Spines” button (placed each spine into its own collection, naming both meshes after the Slicer. VR-SASE created meshes containing the base and tip of the spine, taking its name, and prepending “endpoints”. To improve visualization, and assist with quality control, an empty mesh is placed at the spine distinguishing them from manually segmented dendritic spines.

To obtain surface area metrics, a copy of the original dendrite is created and made hollow with the “Solidify” modifier. Its dendritic spines were removed as before and selected as before.

Next the “Segment Hollow Spines” button is pressed, adding the surfaces to the collection with their corresponding solid spine, prepending “surface” to their name.

### Manual Segmentation

We created a tool to address cases when the VR-SASE Blender addon was unable to segment dendritic spines. To use Manual Segmentation, change Blender’s mode to Edit Mode and select the spine tip by clicking its vertex or face. Pressing the “Manual Segmentation” button places the dendritic spine mesh into a collection containing its endpoints.

If neither automatic or manual segmentation completes successfully, Blender’s native functions suffice to complete the process. In these cases, the dendritic spine was duplicated, “surface_”. prepended to its name, and the faces of its base were removed manually. When endpoints were not positioned correctly, vertices were placed manually marking the dendritic spine base and tip and then joined into a single mesh whose name starts with “endpoints_”. These strings are used to apply different logic to different object categories when the NWB file is created.

### NWB File Generation

After all dendritic spines are segmented, data was written to an NWB file using the “Write NWB File” button. The VR-SASE addon iterates through all the collections in Blender’s “Scene Collection”. For each collection, our addon calculated dendritic spine length based on its endpoints and Blender’s Bmesh programming module determined values for volume, surface area, and center of mass. These parameters along with the metadata from the study were written to an NWB file using the PyNWB library. From the PyNWB library, we used OpticalChannel and ImageSegmentation packages to save the NWB-specified metadata. The physical parameters generated by the VR-SASE Blender addon were added as columns to *plane segmentations*, a structure provided by NWB that united data for each dendritic spine.

### DataJoint Integration

DataJoint is a scientific database framework streamlined for research applications (https://github.com/datajoint). We employed it for the following purposes: 1) Streamlining data integration, management, and analysis. 2) Providing metrics for the Sholl’s analysis by calculating the distance between each spine and the starting point of the dendrite. 3) Refining dendritic spine classification by restricting morphological parameters to published criteria. 4) Ensuring that each dendritic spine has values for length, volume, and surface area.

The VR-SASE Blender addon applied DataJoint functionality by establishing a connection with a remote database using values for host, user_id, and password provided via its GUI This connection allows VR-SASE to use DataJoint completeness’s criteria to ensure data segmentation completion criteria: e.g each spine possesses values for length, surface area, volume and center of mass.

We designed a customized database schema using DataJoint’s schema definition language. where tables represented experimental entities: such as animals, sessions, dendrite morphology, and dendritic spine parameters. The dendrite morphological data needed for the Sholl’s analysis is not part of the NWB data model. We tabulated this data in a csv file which we integrated with the VR-SASE Blender addon. DataJoint ensured that this data was appropriately matched with spine morphology, and NWB metadata. The final table in our pipeline is a Computed table, a class of DataJoint table with extra integrity protection, which calculated the distance between each spine and the starting point of its dendrite, supplying the parameter necessary for the Sholl’s analysis.

Powerful filtering tools enhance DataJoint analyses. Using the DataJoint query language, we constructed queries i.e., filters creating data subsets based on published morphological parameters of dendritic spines. Upper and lower bounds for dendritic spine volume were established as 0.01 μm for small thin spines to 0.8 μm^36, 37^ for large mushroom spines. 3 µm is established as a maximum length for dendritic spines on motor neurons, so we used this criterion^38, 39^. Peters and Kaiserman-Abramof established a 0.5 μm minimum length for cortical spines ^28^, however Harris found dendritic spines as short as 0.2 microns in the hippocampal region. ^36, 37^ There is currently no consensus regarding the minimum length of dendritic spines on motor neurons, so we used Harris’s 0.2um due to the abundance of very small protrusions.

The following query demonstrates application of those parameters and generated our dataset:

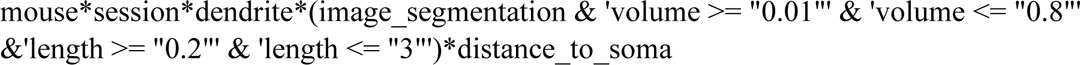

This resulted in data subset containing dendritic spines whose volume was between 0.01 and 0.8µm^3^ and whose length ranged between 0.2 to 3µm. DataJoint refined our pool of 858 potential dendritic spines, to 636 dendritic spines. 94 potential spines were excluded for failing to meet minimum size criteria and 24 were excluded for exceeding it (Supplementary Data.) This data subset was exported as a CSV, and further analyzed in Excel with pivot tables.

Finally, DataJoint applied quality control checks to the dendritic spine segmentation. Because DataJoint ensures that data is complete, it identified dendritic spines which were missing values for length, surface area, or volume.

DataJoint integrated diverse data sources, ensured that spine morphology data was complete, refined our classification of dendritic spines, and enabled a Sholl’s analysis by calculating the distance between dendritic spines and the start of their dendrites. VR-SASE enables researchers to deposit their analysis results directly into a database, promoting research best practices. ^25^

### Sholl’s Analysis

To investigate the distribution of dendritic spines relative to the cell body, we employed a Sholl’s analysis, as described previously ^6, 8, 27, 40, 41^. (See Fig. S2). We calculated the average dendritic spine density within the proximal region, 0-30 μm from the starting point of the dendrite. We utilized Blender’s native 3D modeling features to create a vertex, marking the starting point of the dendrite. To facilitate data binning, concentric circles were placed around this point, incremented by 30 μm. We used Blender’s built-in measurement tool to find the length of dendrites encompassed within each circle and tabulated in a csv file along with the coordinates of the dendrite starting point.

Using a computed table, a table class with extra integrity protection, DataJoint calculated the distance between the center of mass of each spine and the starting point of their dendrite using distance function from the Python math library.

### Distance to soma = math.dist(spine center of mass, soma contact point)

To integrate the measured dendrite length data with the spine morphology data and enable further analysis, the Blender VR-SASE addon linked the .csv file containing the dendrite starting point coordinates and dendrite length measurements to the DataJoint Pipeline, where it was associated with the corresponding spine morphology data, and NWB metadata. The parameter “distance to soma” identified dendritic spines within the circular bins. Density was calculated as follow:

### Spine density = number of dendritic spines in bin/dendrite length

By combining the functionalities of Blender and DataJoint, our Pipeline for data integration and analysis, we established a comprehensive workflow for quantifying dendritic spine density and conducting Sholl‘s analysis. The integration of these tools facilitated accurate and efficient assessment of dendritic spine distribution and provided valuable metrics for our study.

### Quantification and Statistical Analysis

We analyzed tissue from the following groups: SCI and vehicle treatment (n = 5), and sham injured (n = 3). This resulted in 20 dendrites in the SCI group, and 9 in the sham group. This translated to 858 total dendritic spines: 164 dendritic spines in the Sham group and 694 in the SCI group.

We conducted statistical analyses using appropriate tests and evaluated the results based on a significance level of 0.05. We used two-tailed analyses and chose either parametric or non-parametric tests based on the nature of the data. We exported the dataset from the DataJoint pipeline and transferred this data to Prism to perform unpaired t-Tests, or Mann-Whitney Tests as appropriate. All graphs are plotted as mean ± SEM.

## Figure Generation

Figure 1 was created in Biorender.

Figure 2: Neurolucida, ImageJ, and VR-SASE. We generated images for Fig. 2A, B by taking screenshots within the Neurolucida environment (https://www.mbfbioscience.com/products/neurolucida-360/). We processed them with the automated Brightness/Contrast tool in ImageJ to enhance visibility. Fig 2D depicts the maximum size of a dendritic spine in ImageJ. Compression artifacts are large for such a small image, so we used a screenshot to capture a visual representation. Fig. 2E,F were created using Blender’s rendering engine. A virtual camera and lights were placed and positioned around the model in Blender’s 3D viewport, ensuring appropriate visibility of features. Scale was indicated by creating a scale bar in the same plane as the neural tissue.

Figure 3: We analyzed a subset of the Kauer image data using ImageJ to create 3D models, which we initially segmented in Open Brush and then exported to Blender to finish segmentation. Blender created NWB files for each dendrite, which are in the linked DANDIset. A DataJoint database pipeline refined the dataset based on morphological parameters and calculated the distance between each spine and the start of its dendrite. The filtered DataJoint pipeline was exported as a CSV, which served as the data source for pivot tables in V1.5_DiscDataCompilation. Graphs and statistics were created in Prizm.

Figure 4: We reconstructed a neuron from the DIADEM challenge. We used ImageJ, Open Brush, and Blender as above and created a 2D image of our reconstruction with Blender’s rendering tool (OP-09_VR-SASE.png) Fig 4B. The HBP Neuron Morphology Viewer transformed the DIADEM reconstruction into 2D format (OP-09_GoldStandard.png) Fig 4C. Both images were registered on the maximum projection of the original neuron in Photoshop (OP-09_Both_aligned.psd). The maximum projection (MAX_OP-09.png) Fig. 4A was enlarged to the size of the gold standard, and our reconstruction was scaled down. Each reconstruction super-imposed over the original tissue was exported from Photoshop and regions with the maximum projection visible were traced and measured Fig 4. D, E. (OP-09_VR-SASEgreen_ROI.tif and OP-09_GoldStandardgreen_ROI). Fig 4 F, G are renderings of our segmentation results, randomly colored by Chatgpt. The Blender files contain scripts to adjust colors and their outputs are OP-09_Rainbow and OP-09_RainbowCloseup.

Figure 5: The post hoc analyses were created in Chatgpt and executed in Blender’s scripting environment, prior to rendering the images. Within the Blender file, L712_Dendrite1aMushroomConvexHull.blend, the script “ConvexHull” created a convex shell around the dendritic spine, and its head. Meshes in the collection, MushroomSpineConvexHull were examined through Blender’s UI, and volumes corresponding to the spine, head, and convex hulls are commented in this script. To assist with visualization, the hulls were also converted to wireframes.

The Cord Length Distribution Histogram requires multiple steps to recreate. Within the L712_023-01-31_16.11.24_Dendrite1aMushroomCord.blend, there are 3 scripts. “AddCylinders”, “ColorCylinders”, and “ExportCylinderNames”. The first script creates a csv containing cord names and lengths. Unrealistic cords were removed manually, the remaining cord names using “ExportCylinderNames” and compared to the original list in Excel, which deleted unique values from the original list, matching the remaining cords with their length (MushroomCords.csv). The cord lengths were graphed in Prizm. (MushroomCords.pzfx.) (Fig 5C). Figures 5A, B, D, E were created using Blender’s rendering engine as described above.

## Diadem Validation

To generate the image for validating the accuracy of VR-SASE, we obtained raw images and the gold standard SWC file for OP-09 from the olfactory projection fiber dataset from the DIADEM challenge: https://diadem.janelia.org/olfactory_projection_fibers_readme.html We used the Neuroinformatics Morphology Viewer to convert the swc file into a png file: https://neuroinformatics.nl/HBP/morphology-viewer/<u>#</u>. We then created a maximum projection of OP-09 using ImageJ’s Z project function to create a flattened representation of the stack of raw image, which we saved in png format.

We transferred both the gold standard reconstruction and maximum projection images to Photoshop, creating a layer for each. We then scaled up the maximum projection so that it aligned with the gold standard reconstruction. Our reconstruction, rendered in Blender as described above, was added as an additional layer in Photoshop, then scaled down to the size of the gold standard, and aligned with the maximum projection. Both super-imposed reconstructions were flattened and exported as flattened Tiff files with 3000 x1500 pixels for analysis in ImageJ, where they were scaled to 168.78 x 84.39 microns. Regions where the maximum projection was visible represent modelling inaccuracies and were quantified by tracing regions of interest (ROIs) with ImageJ’s polygon selection tool, enabling precise quantification.

### Post Hoc Analyses

#### Cord Length Distribution Histogram

CLDH facilitates partitioning morphologically similar dendritic spines into clusters, further refining analyses of dendritic spine morphology^16^. A CLDH is a probability distribution of the lengths of internal cords connecting vertices of a dendritic spine mesh. To perform a post-hoc CLDH analysis, we used the Text Editor (See Figure S2), Blender’s native python scripting environment to create a set of internal cords for a representative dendritic spine. For each vertex, we created cords for 1 in every ten of their possible connections, excluding vertices within 0.2 µm. This created a rich, but manageable set of cords. Our post-hoc analysis also created a csv file containing the name and length of each cord. Cords extending over concave surfaces represent unrealistic geometries and were removed manually. The lengths of the remaining cords are the basis for our CLDH.

### Convex Volume Index

CVI is based on the difference in volume between a dendritic spine and its convex hull. The convex hull is the smallest possible ‘shell’ enclosing all vertices of the dendritic spine head mesh without having any concave surfaces.^12^ To perform a *post-hoc* CVI calculation, we used a python script executed in the Text Editor to create a convex hull surrounding the dendritic spine head. We obtained the volume of the convex hull from the 3D Printing addon and compared it to the volume of the dendritic spine.

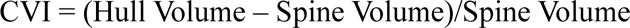

We next extended the methodology proposed by Kashiwagi et al to calculate the ratio for the entire dendritic spin.

## Hardware

### Segmentation was conducted on multiple platforms, including

Alienware Desktop PC with Intel® Core™ i7-10700k CPU @3.80 GHz with 16.0 GB installed RAM running Windows 11 Home Operating System 22H2 with the Oculus Rift S VR headset, from Meta Platforms.

Alienware Laptop PC with 12^th^ Gen Intel® Core™ i9-12900HK, 2500 MHz, 14 Cores, Logical Processors 32.0 GB installed RAM running Windows 11 Home Operating System tethered to an Oculus Quest 2 VR headset, from Meta Platforms.

Blender Segmentation was carried out on these machines, as well as computers with the following specifications:

- Intel(R) Core(TM) i7-1065G7 CPU @ 1.30GHz with 32.0 GB installed Ram running Windows 11 Home Operating System 22H2.
- 16-inch MacBook Pro Apple M2 Max with 12-core CPU, 38-core GPU, 16-core Neural Engine 96GB unified memory MacOS 13 Ventura.

### Large Language Model Usage

Chat GPT-3 generated code for the post-hoc analyses. We employed it during programming for prototyping and troubleshooting and during the early stages of manuscript drafting, and to create the color scheme used to display segmentation results from the DIADEM Challenge.

After using Chat GPT-3, the authors reviewed and edited the content as needed and take full responsibility for the content of the publication.

### Supplemental Items

VR-SASE Supplemental Information VR-SASE User documentation VR-SASE Video Overview

## Resource Availability

### Lead Contact

Further information and requests for resources and reagents should be directed to and will be fulfilled by the lead contact, Andrew Tan.

## Materials Availability

This study did not generate new unique reagents.

## Data and Code Availability

Neurodata Without Borders data will be deposited at Dandi Archives and are publicly available as of the date of publication. Accession numbers will be listed in the key resources table.

Images, Blender Files, NWB files, and analysis products have been deposited at Data Dryad will be publicly available as of the date of publication. DOIs are listed in the key resources table.

Blender addon code is currently in the following Github repository: https://github.com/ycnrr/VR-SASE_Blender-Addon

DataJoint Code and queries are currently in the following Github repository: https://github.com/ycnrr/Spasticity-DataJoint

Upon publication, code will be made publicly available. DOIs will be listed in the key resources table.

Adjective/all data reported in this paper will be shared by the lead contact upon request.

## Acknowledgements

The work is funded by grants from the PVA, VA Medical and Rehabilitation Research Services, The Taylor Foundation, the Erythromelalgia Association, and the Flaherty and Wilshire Walsh Endowments, as well as the SPiRE Program. It is conducted in collaboration between the Paralyzed Veterans of America and Yale University at the Center for Neuroscience and Regeneration Research.

