## Supplemental for "A Novel, Open-Source Virtual Reality Platform for Dendritic Spine Analysis"

### VR-SASE Interface

The VR-SASE workflow begins with model creation from neural images in ImageJ. Laborious tracing methods are replaced by positioning discs on room-scale VR neural models, efficiently partitioning the neuroanatomy. Our custom addon for the 3D modelling tool Blender uses these discs to segment the dendritic spines. To promote data management best practices, VR-SASE automates data extraction and integrates data with NWB and DataJoint. This emerging best practice in data handling within the field of neuroscience ensures that the data produced is in a standardized and easily shareable format, promoting transparency and reproducibility in research

The VR-SASE Blender Addon user interface (UI) is shown in **Fig. Supplemental 1. Fig. S1A** shows a segmented neuron alongside the VR-SASE UI. The fields necessary to provide complete, experiment-level metadata for NWB are shown in **Fig. S1B.** Tools we developed to streamline segmentation are shown in **Fig. S1C. Fig. S1D** provides the automated interface to write morphological data to NWB files, including dendritic spine density, length, volume, and surface area. These measures play a crucial role in understanding synaptic strength [1], electrical interactions[2, 3], and calcium dynamics[4]. **Fig. S1E** depicts the interface that integrates VR-SASE with DataJoint for quality control and advanced analytics. Providing the host, user ID, and password enables VR-SASE to integrate with a database, a best practice in scientific data management[5]. **Fig. S1F** shows the result of Segmentation in Blender’s scene collection: A spine mesh, its endpoints, and its surface are placed into a collection.

### VR-SASE UI

The “Separate Meshes” button on the VR-SASE Segmentation Tools Panel (**Fig. S1C**) creates individual meshes of each spine.

The separated spines must have their origin updated through Blender’s Graphical User Interface (GUI). Then the ‘Slicer’ discs must be placed in a collection. To automatically segment the spines, they were selected along with the collection of slicers as shown in (**Fig. S1F**).

The “Segment Solid Spines” button (**Fig. S1C**) placed each spine into its own collection, each taking the name of the Slicer. VR-SASE created meshes containing the base and tip of the spine, taking its name, and prepending “endpoints”. To improve visualization, and assist with quality control, an empty mesh is placed at the spine distinguishing them from manually segmented spines.

To obtain surface area metrics, a copy of the original dendrite is created and made hollow with the “Solidify” modifier. Its spines were removed as before and selected as before. Next the “Segment Hollow Spines” button (**Fig. S1C**) is pressed, which adding them to the collection with their corresponding solid spine prepending “surface” to their name.

### Stubby Spines

Stubby spines represented a challenge because their variable morphology requires multiple measures and rules to disambiguate.[6] **Fig. Supplemental 3** depicts the challenges of quantifying the morphology of stubby spines. For all spines that are not stubby, the tip is defined as the farthest point from the base. **Fig. S3A** shows that placing the spine tip at the location farthest from the spine base does not reflect the true tip. We used Blender’s raycasting functionality to determine the endpoints of Stubby Spines. The raycast begins at the base of the spine, extending in the direction of the spine’s center of mass, and terminates when it reaches the surface of the spine. **Fig. S3B** demonstrates the utility of raycasting in a cone instead of a single line, which can place the spine tip correctly. We found that the farthest point from the spine base, when restricted to a raycast cone with a 15° angle, performed more reliably than a simple line raycast. Nevertheless, this method requires researchers to inspect and correct the endpoints of the spine. We manually corrected the endpoints of stubby spines when they were placed on the rim of the spine base.

### Supplemental Figure Descriptions

**Fig. Supplemental 1**: VR-SASE User Interface: **Fig. S1A** shows a segmented neuron. **Fig. S1B** shows the fields that provide NWB’s prescribed metadata. **Fig. S1C** shows the tools we developed to streamline segmentation: Separate Meshes, Segment Solid Spines, Segment Hollow Spines, and Manual Segmentation. **Fig. S1D** depicts the interface that automates data extraction and writes NWB files. **Fig. S1E** depicts the interface that integrates VR-SASE with DataJoint for quality control and advanced analytics. **Fig. S1F** shows the result of Segmentation in Blender’s scene collection: A spine mesh, its endpoints, and its surface are placed into a collection.

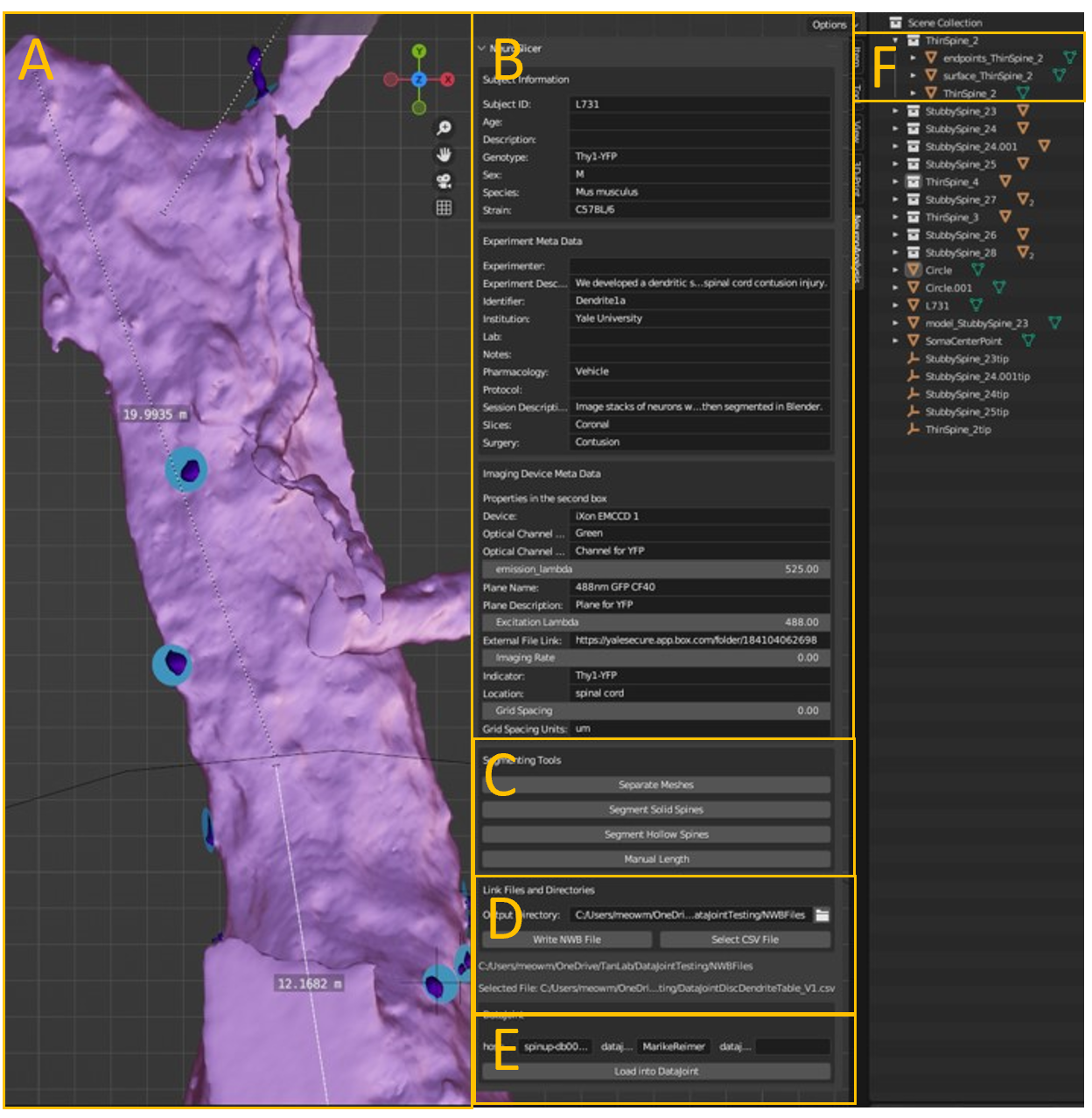

**Fig. Supplemental 2: Sholl’s Analysis**. Orange circles, 30 µm in diameter, originate from the starting point of the dendrite. Dendrite length was measured with Blender’s native Grease Pencil on an orthogonal view to the neural tissue. The number of dendritic spines per unit of dendrite length determined its density. The center of mass of the dendritic spine and the starting point of its dendrite was used to bin the spines according to their distance from it.

The panel beneath the Sholl’s analysis shows Blender’s Text Editor, an interface for rapid prototyping/post processing with the Python programming language.

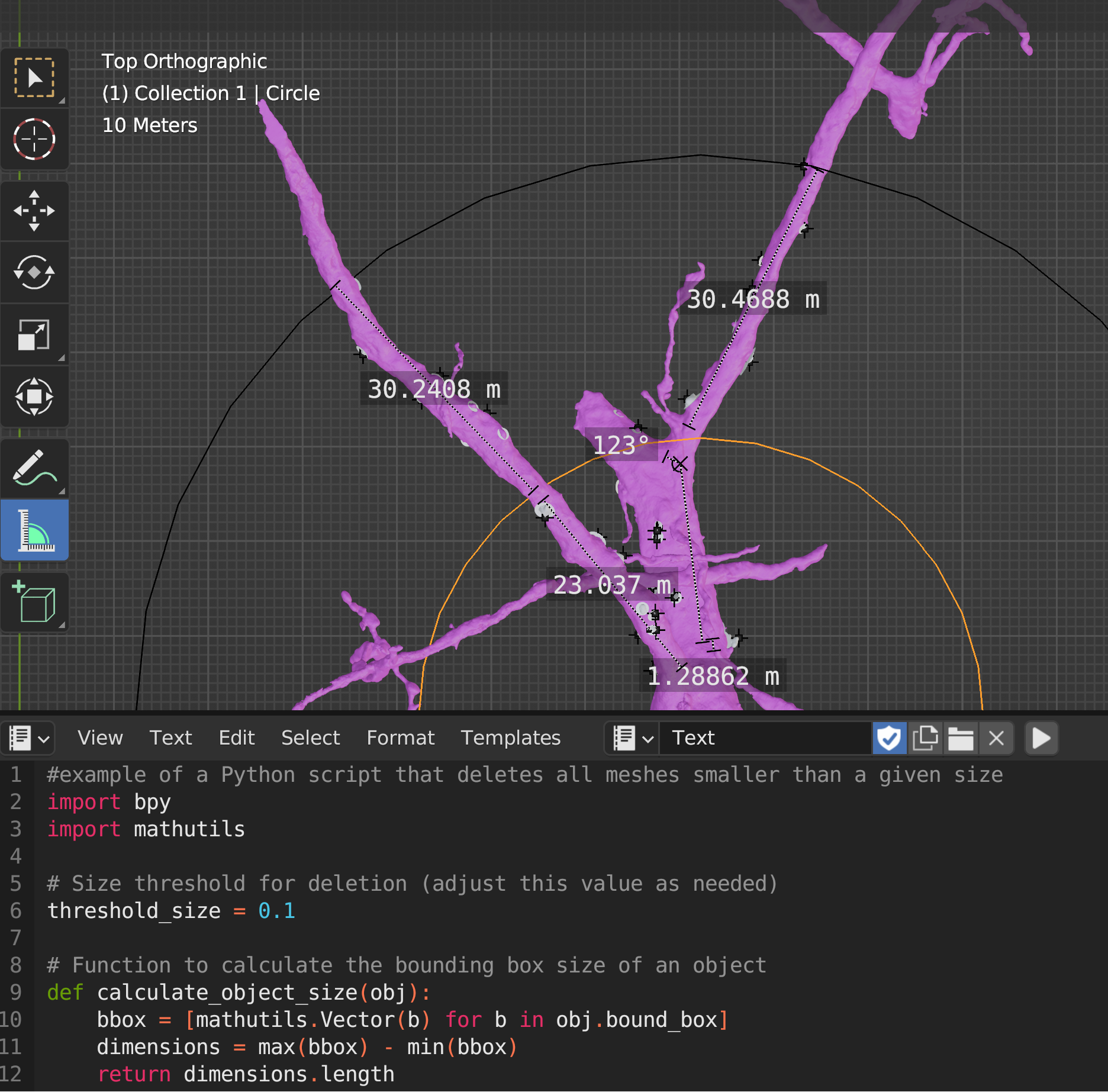

**Fig. Supplemental 3: Stubby Spine Morphology**. One challenge in quantifying the morphology of stubby spines lies in the ambiguity of length. **Fig. S3A** shows that placing the spine tip at the location farthest from the spine base does not reflect the true tip and the raycasting method we used to mitigate this issue. Raycasting is Blender functionality that projects a line from a specified point, in a specified direction that proceeds until it encounters an object. **Fig. 3B** demonstrates the advantage of raycasting in a cone instead of a single line.

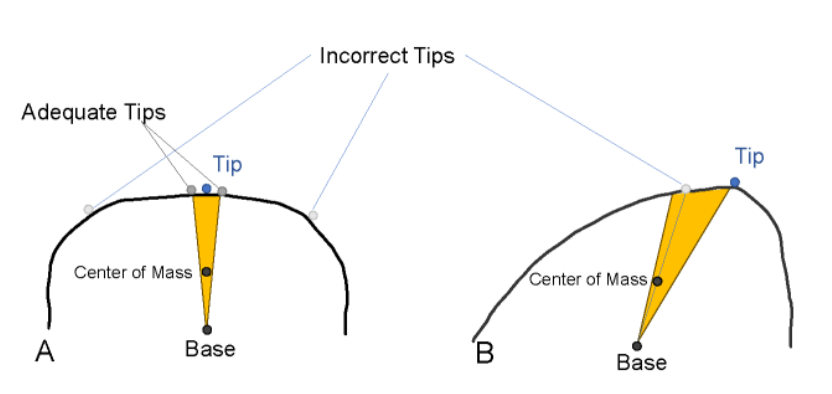
